# Direct recruitment of RNA Polymerase II by NANOG to activated target genes

**DOI:** 10.1101/2025.11.27.690710

**Authors:** Nicholas Paul Mullin, Elisa Barbieri, Jingchao Zhang, Matúš Vojtek, Alison McGarvey, Douglas Colby, Simon Tomlinson, Ian Paul Chambers

**Affiliations:** Centre for Regenerative Medicine, Institute for Regeneration and Repair, University of Edinburgh, 5 Little France Drive, Edinburgh EH16 4UU, Scotland; Institute for Stem Cell Research, School of Biological Sciences, University of Edinburgh, 5 Little France Drive, Edinburgh EH16 4UU, Scotland

## Abstract

Cell identity relies upon transcription factors (TFs). The concentration of the TF NANOG determines the efficiency of maintenance of mouse embryonic stem cell (ESC) identity. However, the mechanisms by which NANOG acts are not fully understood. Models of mammalian transcription generally propose that TFs bind DNA and connect to the transcriptional machinery indirectly via intermediary proteins. Accordingly, examples of direct contact between cell-type specific TFs and RNA synthesis enzymes in mammalian cells remain elusive. Here we show that NANOG directly contacts RNA Polymerase II (RNAPII) via aromatic residues within the low complexity domains of both proteins. NANOG can localize RNAPII to a specific DNA site, with RNAPII enhancing NANOG DNA affinity. The NANOG-RNAPII complex is dissociated by the transcriptional pause-release enzyme, CDK9. Inhibition of CDK9 enhances NANOG chromatin binding, while induction of NANOG localizes RNAPII to specific NANOG chromatin sites. Induced RNAPII localization is selective for targets activated by NANOG. A NANOG mutant that retains DNA, but not RNAPII binding, cannot stimulate RNAPII localization to chromatin, shows an impaired transcriptional response and does not drive LIF-independent ESC self-renewal. These results identify a novel, regulatable interaction between a cell-type specific TF and RNAPII that can change transcription and alter cell fate.

## Introduction

Self-renewal of mouse embryonic stem cells requires the action of extracellular signals, including LIF (*1, 2*) as well as the action of intrinsic transcriptional regulators centred around the transcription factors (TFs) SOX2, OCT4 and NANOG (*3–8*). Of these, NANOG is able to circumvent the need for LIF when overexpressed (*7, 8*). Indeed, the concentration of NANOG determines the efficiency of maintenance of ESC self-renewal (*7–9*). Although prior work has identified NANOG target genes (*10*) and partner proteins (*11*), the mechanisms by which NANOG effects efficient ESC self-renewal are not fully understood.

The primary sequence of NANOG indicates the presence of a homeodomain DNA binding region embedded in sequence devoid of additional recognizable protein domain sequence. The most eye-catching sequence is a low complexity domain (LCD) consisting of 10 imperfect repeats in which every 5th residue is a tryptophan. This LCD is commonly referred to as the tryptophan repeat (WR) and is required for NANOG to confer LIF-independent ESC self-renewal (*12–14*).

In this work, we have pursued the identification of RNA Polymerase II (RNAPII) within the NANOG interactome (*11*). This shows that NANOG binds directly to the C-terminal domain of RNAPII. Our findings have implications for the connectivity of TFs to the basic transcriptional machinery, as well as for the actions subsequently initiated at enhancers by TF binding.

## Results

### A NANOG binding motif recurs in NANOG interacting proteins

Our previous analysis identified an interaction between NANOG and SOX2 in which tyrosines within a motif (SX[S/T]Y) in SOX2 bind directly to tryptophans of the NANOG WR (*11*): (Fig. 1a gives an overview of the primary NANOG structure). Analysis of the NANOG interactome (*11*) for additional proteins containing an SX[S/T]Y motif identified 35 proteins, 8 of which have >1 SX[S/T]Y motif (Supplementary Data Table 1). Of these, RNAPII is unique, as it alone contains multiple, consecutive consensus repeats. Specifically, the RNAPII CTD contains 33 copies of the SX[S/T]Y motif (Fig. 1b). In mammals, the CTD is composed of 52 repeats of the consensus heptad (Y^1^S^2^P^3^T^4^S^5^P^6^S^7^) that functions as a minimal binary unit (*15*). The heptad is an evolutionary ancient sequence, proposed to have arisen by conjunction of two motifs, one of which is SP[S/T]Y (*16*).

**Figure 1.**
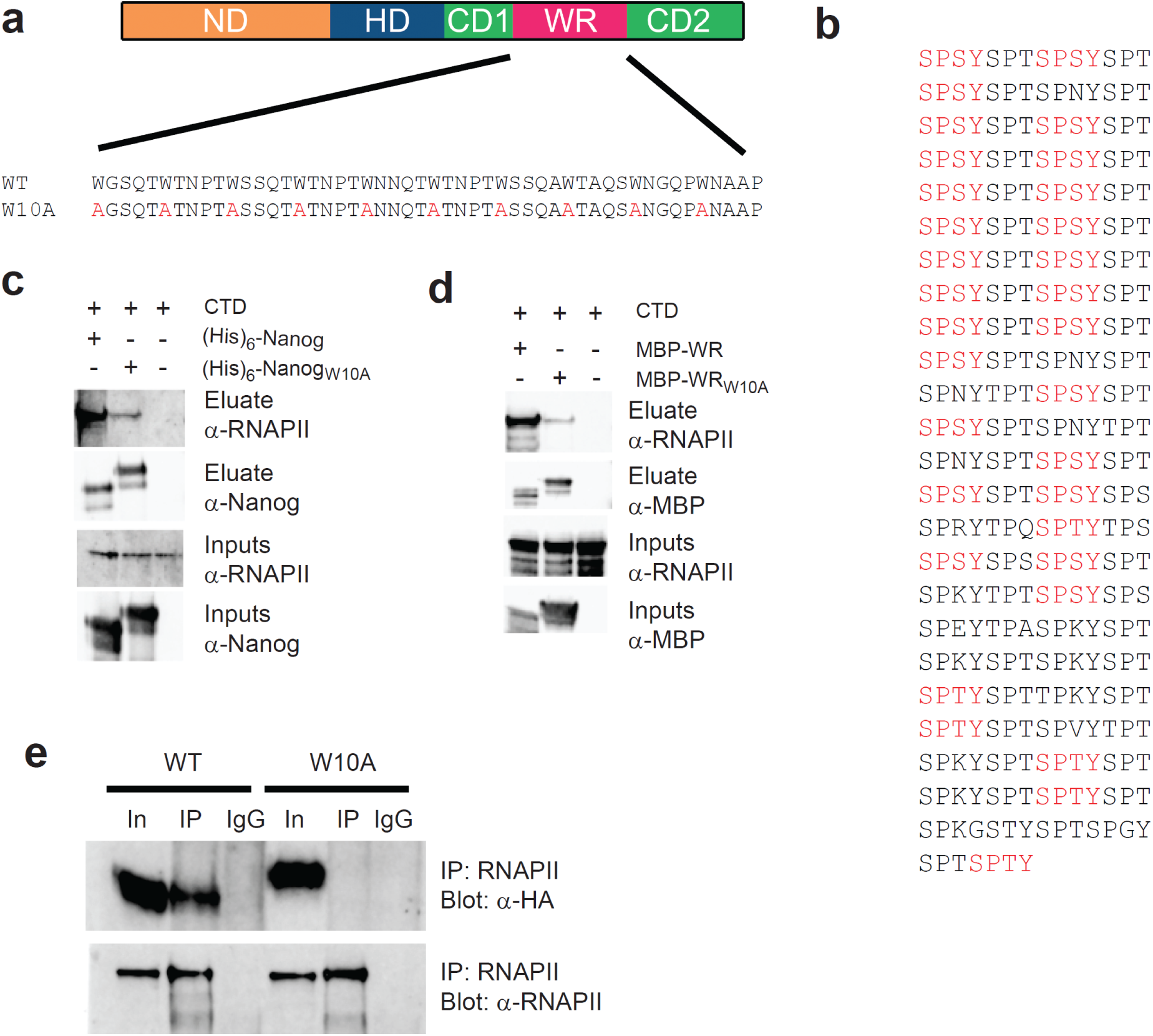
The NANOG WR interacts directly with the RNA polymerase II CTD. **a.** Top - Domain structure of NANOG. ND – N-terminal domain, HD – homeodomain, CD1 – C-terminal domain 1, WR – tryptophan repeat, CD2 – C-terminal domain 2. Bottom - Sequences of WR of wild-type (WT) and W10A mutant. **b.** The amino acid sequence of the CTD of mouse RNAPII II. All copies of the Sx[S/T]Y sequence identified in SOX2 are highlighted in red. **c.** GST-CTD was co-expressed in *E. coli* alongside either His-tagged NANOG or His-tagged NANOG_W10A_ and NANOG purified on nickel resin. Co-purifying GST-CTD was detected by immunoblotting with anti-RNAPII (n = 2). **d.** MBP-WR or MBP-WR_W10A_ were co-expressed with GST-CTD in *E. coli*. MBP-NANOG variants were purified on amylose resin and analysed for co-purifying GST-CTD by immunoblotting with anti-RNAPII (n = 2). **e**. Co-immunoprecipitation of NANOG or NANOG-W10A and RNAPII from ESCs (n = >10).

### Direct binding of NANOG to RNAPII

The above observations raise the hypothesis that RNAPII may bind NANOG directly through interaction of RNAPII CTD tyrosine residues with NANOG WR tryptophan residues. To test this hypothesis, full length His-tagged-NANOG was co-expressed alongside RNAPII CTD in *E. coli* to enable assessment of protein interactions in the absence of mammalian post-translational modifications. NANOG was purified from lysates on nickel-agarose and co-bound protein detected by immunoblotting (Fig. 1c). NANOG retained RNAPII. In contrast, binding of RNAPII to a mutant NANOG in which the WR tryptophans were substituted by alanines (NANOG_W10A_) (Fig. 1a) (*13*) was strongly reduced (Fig. 1c). This establishes that NANOG binds RNAPII directly and indicates that WR tryptophans are required for efficient RNAPII binding.

To determine whether the WR is sufficient to bind RNAPII, the WR (or the W10A mutant version) was isolated from the rest of NANOG and expressed as a MBP-fusion alongside CTD. Purification of MBP-complexes showed that WR bound RNAPII and confirmed that binding of WR_W10A_ to RNAPII was strongly reduced (Fig. 1d). Therefore, the low complexity domains of NANOG and RNAPII are sufficient for interaction between the two proteins in vitro, with tryptophan residues in the WR fulfilling a critical role. To determine whether NANOG and RNAPII also interact in vivo, RNAPII was immunoprecipitated from ESCs. Analysis of co-precipitating proteins showed that NANOG was found in RNAPII precipitates but NANOG_W10A_ was not (Fig. 1e). These results show that NANOG directly contacts RNA Polymerase II (RNAPII) via aromatic residues within the low complexity domains of both proteins.

### NANOG acts as a bridge between a specific DNA site and RNAPII

The above results also suggest that binding to WR may be used to localize RNAP2 to a specific DNA recognition site. To assess this, the interaction of NANOG with RNAPII was further investigated using an electrophoretic mobility shift assay (EMSA). Addition of purified NANOG to a NANOG specific oligonucleotide(*17*) retarded DNA mobility due to NANOG:DNA complex formation (Fig 2a). Comparative titration of NANOG and NANOG_W10A_ showed no difference in the affinity of NANOG and NANOG_W10A_ proteins for DNA (Supplementary Data Fig. 1). Interestingly, while purified CTD did not bind DNA on its own, addition of CTD to DNA in the presence of NANOG altered the mobility of the labelled DNA, indicating formation of a ternary DNA:NANOG:RNAPII complex (Fig. 2b). In contrast, CTD addition did not affect the mobility of the NANOG_W10A_:DNA complex (Fig. 2b). These results indicate that NANOG can directly connect a NANOG-specific recognition site on DNA to RNAPII and that this connection is dependent upon WR tryptophans. Furthermore, titrating NANOG into the EMSA in the presence or absence of a fixed concentration of CTD showed that the CTD enhances the affinity of NANOG for DNA approximately 2-fold (59.2 ±5.3nM vs 34.6 ±3.6nM) (Fig. 2c).

**Figure 2.**
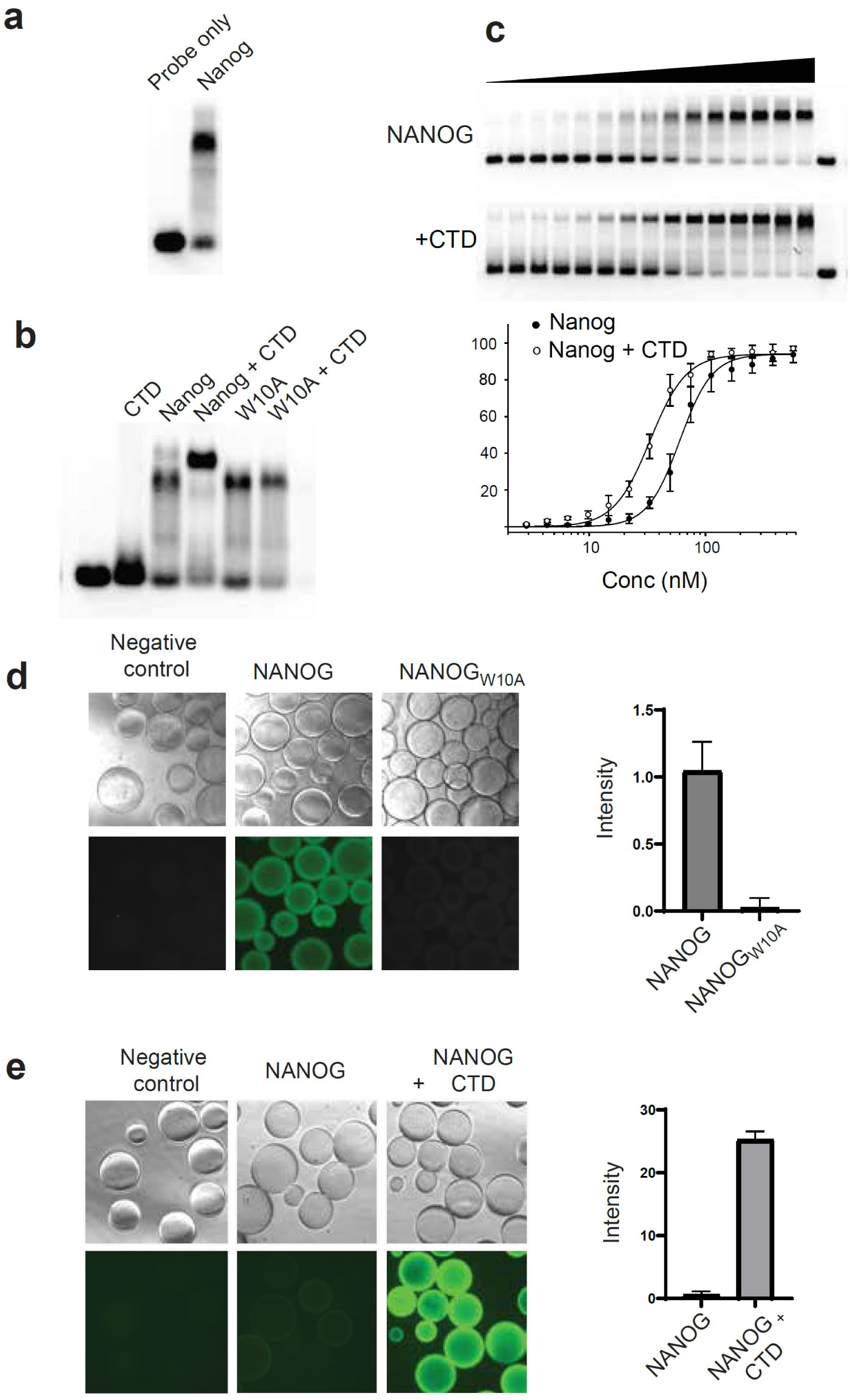
The interaction of NANOG with DNA is enhanced by the CTD. **a, b.** A fluorescently tagged 26mer oligonucleotide was incubated with the indicated proteins and assessed by EMSA (n =3). **c.** EMSA with NANOG titrations performed in the absence (top) or presence (middle) of 1.5μM CTD. NANOG was titrated from 500nM in a 1/3 dilution series. The percent shifted probe was quantitated (bottom) (n=3, error bars represent standard deviation). **d**. NANOG bridges between DNA molecules as shown by BASIL (Supplementary Data Fig. 2). DNA was bound to a bead surface via a 5’ biotin. After incubation with NANOG or NANOG_W10A_, a fluorescent oligonucleotide will only bind to immobilized NANOG if it exists as a multimer (n=3). Right, normalized mean fluorescence after subtraction of the intensity of the negative control; error bars are standard deviation. Quantitation was performed using Cellpose. **e**. BASIL assay in the presence of CTD. After immobilization of DNA to the bead surface, beads were incubated with NANOG or NANOG plus CTD. Quantitation was performed as in d.

The ability of CTD to augment NANOG binding to DNA is further demonstrated by an assay in which fluorescently labelled oligonucleotide is localized to a bead surface, in a **B**ead **As**say of **I**nteracting **L**igands (BASIL) (Supplementary Data Fig. 2). NANOG forms dimers (*12, 14*) which may allow NANOG to bridge two DNA binding sites. To test this, a biotinylated oligonucleotide containing a single NANOG binding site is bound to streptavidin beads before incubation with NANOG. Subunits of a NANOG oligomer not bound directly to DNA may then be available to bind further DNA molecules. If so, subsequent incubation of beads with a fluorescently labeled oligonucleotide containing a NANOG binding site would localize fluorescence to the bead surface. Incubation of NANOG in this assay localizes fluorescence to the bead surface. This did not occur with NANOG_W10A_ (Fig. 2d) consistent with NANOG_W10A_ being monomeric(*13*). Intriguingly, when the RNAPII CTD is incubated with NANOG, fluorescence is markedly enhanced (Fig. 2e), suggesting that the multivalent CTD may recruit additional NANOG subunits. Together, these results show that NANOG can localize RNAPII to a specific DNA site, with RNAPII enhancing NANOG DNA binding.

### The WR mediates transcriptional responses to NANOG binding

The preceding results raise the possibility that WR binding of RNAP2 may contribute to gene regulation by NANOG. Previous work has identified a cohort of ~100 NANOG-responsive genes(*10*) out of the several thousand sites in ESC chromatin that bind NANOG (*18–20*). To assess the effect of mutation of WR tryptophans on regulation of gene transcription by NANOG, we treated *Nanog*^−/−^ ESCs expressing NANOG-ERT^2^ (*10*) or NANOG_W10A_-ERT^2^ fusions with tamoxifen and measured changes in gene expression over a six hour time course. Changes in gene expression in response to NANOG activity were either absent or strongly diminished in response to NANOG_W10A_ (Fig. 3a).

**Figure 3.**
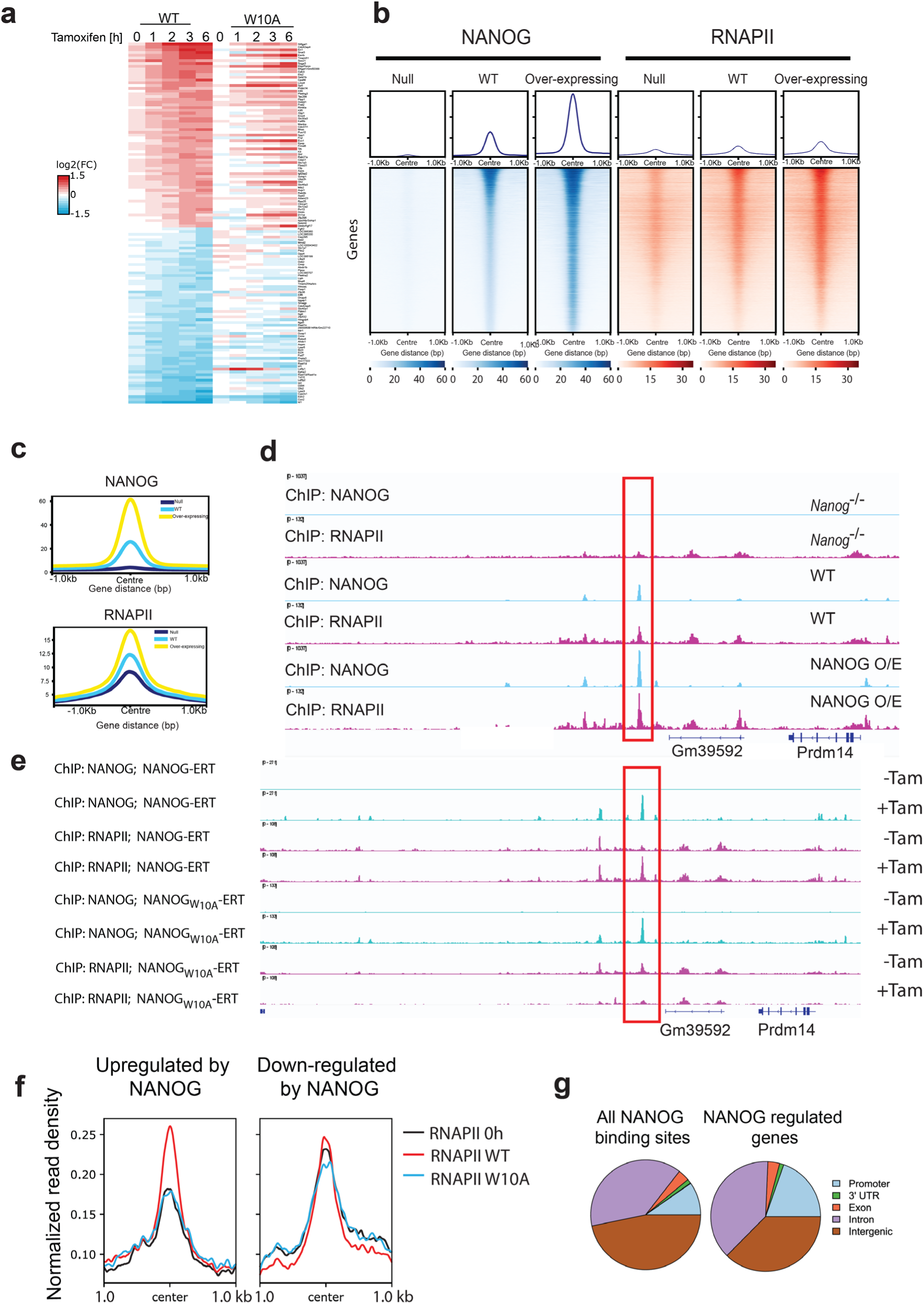
Nanog alters transcription and DNA binding by RNAPII. **a.** RNA timecourse of NANOG-ERT^2^ and NANOG_W10A_-ERT^2^ cells treated with tamoxifen as indicated. Genes which are up- or down-regulated are shown. **b.** ChIP-seq heatmap showing NANOG and RNAPII occupancy in *Nanog*^−/−^, WT and NANOG over-expressing ESCs. **c.** Average NANOG and RNAPII ChIP-seq signal in *Nanog*^−/−^, WT and NANOG over-expressing ESCs **d**. ChIP-seq tracks for NANOG and RNAPII around the *Prdm14* locus in *Nanog*^−/−^, WT and NANOG over-expressing ESCs. The red box highlights where increased NANOG signal results in increased RNAPII binding. **e.** ChIP-seq tracks for NANOG-ERT^2^, NANOG_W10A_-ERT^2^ and RNAPII at the *Prdm14* locus with (+) or without (-) addition of tamoxifen. The red box highlights the RNAPII peaks increased by the presence of NANOG. **f.** Average RNAPII ChIP-seq signal at putative regulatory elements associated with positively and negatively regulated NANOG target genes before or after induction of NANOG-ERT^2^ and NANOG_W10A_-ERT^2^. **g**. Venn diagrams showing the binding of NANOG to sites in the genome and at genes regulated by NANOG.

### The WR recruits RNAPII to upregulated target genes

To investigate whether NANOG influences chromatin localization of RNAPII, ChIP-seq for NANOG and RNAPII was performed in cells expressing a range of NANOG concentrations. Analysis of *Nanog*^−/−^ (*9*), wild-type and NANOG over-expressing ESCs (*21*) demonstrates that as the NANOG level increases, the amount of RNAPII that co-localizes at NANOG-bound chromatin sites also increases (Fig. 3b, c, d, Supplementary Data Fig 3). To investigate dynamic changes in RNAPII recruitment to chromatin, ChIP-seq for NANOG and Poll II was performed in cells expressing NANOG-ERT^2^ or NANOG_W10A_-ERT^2^ in the presence or absence of tamoxifen. As expected, tamoxifen induced the localization of NANOG-ERT^2^ and NANOG_W10A_-ERT^2^ to target gene chromatin (Fig. 3e, Supplementary Data Fig. 4a, 5). However, while localization of NANOG-ERT^2^ to putative regulatory elements near NANOG target genes was accompanied by recruitment of RNAPII to overlapping sites at multiple time points (Fig. 3e, Supplementary Data Fig. 4a, 5), localization of NANOG_W10A_-ERT^2^ to chromatin did not stimulate RNAPII recruitment (Fig. 3e, Supplementary Data Fig. 4a, 5). This induced localization of RNAPII was selective for targets activated by NANOG as it did not occur at putative regulatory elements associated with genes repressed by NANOG (Fig. 3f, Supplementary data Fig. 6). NANOG has been shown to be localized globally at sites distal to gene promoters (*6, 22, 23*). Our ChIP-seq analysis confirms and extends this by showing that sites associated with differentially expressed genes (DEGs) are also predominantly distal to promoters (Fig. 3g), irrespective of their association with positively or negatively regulated DEGs (Supplementary Data Fig. 4b). Together these results indicate that WR tryptophans are required for both the recruitment of RNAPII to NANOG responsive loci and for activation of gene transcription in response to NANOG.

### Selective binding of NANOG to phosphorylated forms of RNAPII

The central role of tryptophan residues in the NANOG-RNAPII complex suggests an interaction mediated by stacking of aromatic residues, as proposed for the interaction of SOX2 with NANOG (*11*). Such a hydrophobic interaction may be expected to be sensitive to the introduction of charged residues nearby. The RNAPII CTD can be extensively modified by phosphorylation (*24*). To determine the effect of CTD phosphorylation on the interaction of RNAPII with NANOG, antibodies recognizing phospho-specific forms of RNAPII CTD were used to assay co-immunoprecipitation of NANOG with RNAPII from ESCs. Precipitation of NANOG with antibodies recognizing phosphorylation at, or adjacent to, the CTD tyrosines would not support the hypothesis of interaction via aromatic stacking. However, RNAPII antibodies recognising phospho-serine2 or phospho-tyrosine1 did not co-immunoprecipitate NANOG (Fig. 4a). In contrast, NANOG was co-immunoprecipitated using an anti-RNAPII antibody recognizing phosphorylation of serine-5 (Fig. 4a). These results suggest that introduction of negative charge on, or immediately adjacent to the tyrosine residues of RNAPII-CTD prevents the interaction with NANOG, supporting a model in which NANOG tryptophan residues and RNAPII tyrosine residues interact via aromatic stacking.

**Fig. 4.**
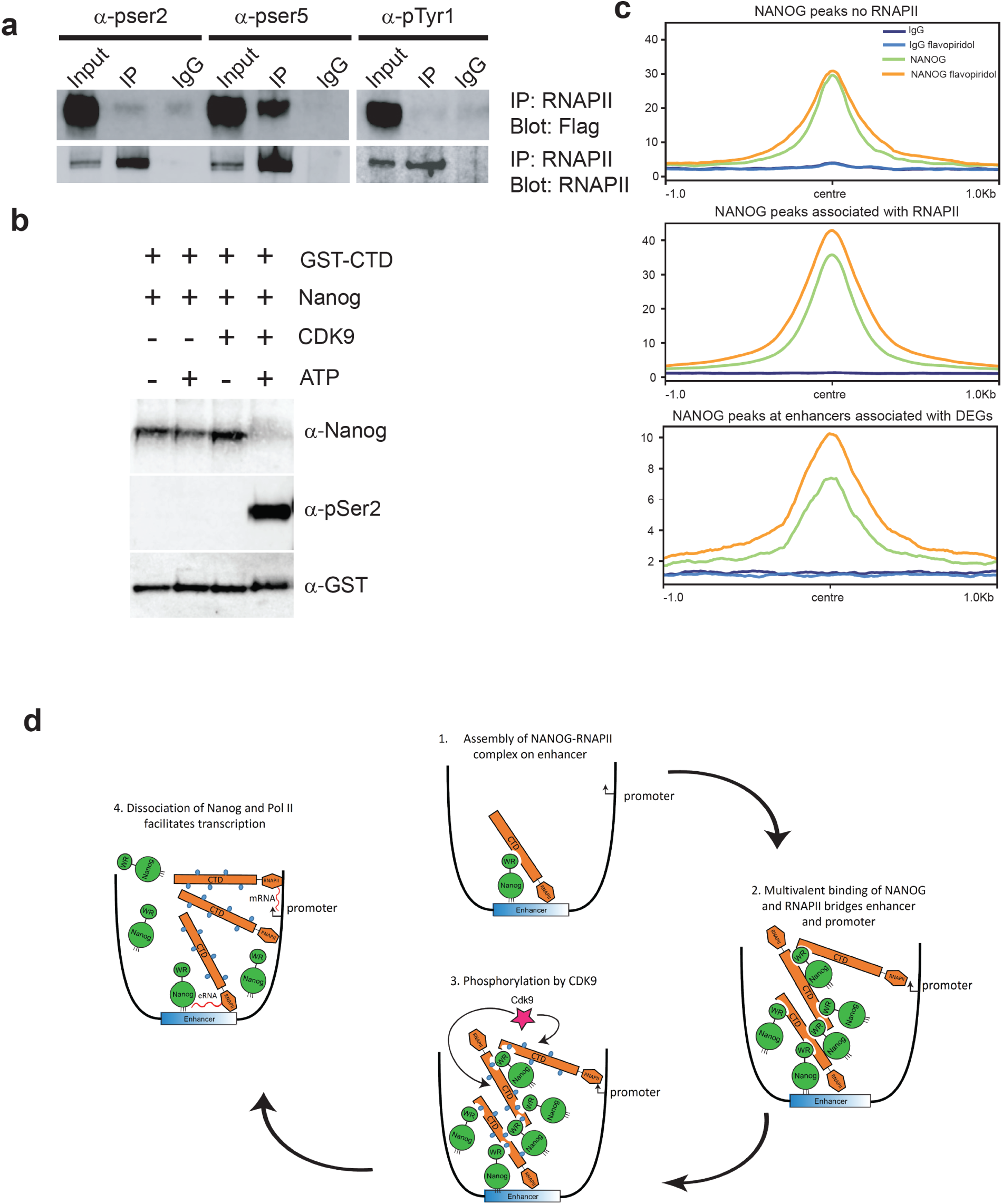
The NANOG – RNAPII interaction is sensitive to phosphorylation. **a.** (Flag)_3_-NANOG was expressed in E14/T ESCs and interaction with RNAPII assessed by immunoprecipitation (IP) with phospho-specific anti-RNAPII antibodies, followed by immunoblotting (IB) as indicated (n = 3). **b.** GST-CTD and NANOG were co-expressed in *E. coli* and purified on GSH resin. The CTD-NANOG complex bound to resin was incubated as indicated and protein remaining bound to the resin analysed by immunoblotting with the indicated antibodies (n = 3). **c**. Inhibition of CDK9 results in accumulation of NANOG at enhancers. E14Tg2a ESCs were treated with flavopiridol and NANOG ChIP performed. Intensity of NANOG binding at peaks with no associated RNAPII, at peaks associated with RNAPII, and at peaks associated with NANOG differentially expressed genes. **d**. Model of NANOG-RNAPII interaction at enhancers. NANOG binds to DNA at an enhancer and recruits RNAPII via WR-CTD binding (1). Multivalency on RNAPII CTD and NANOG WR enables further recruitment of NANOG and RNAPII monomers leading to the growth of a NANOG-RNAPII domain from the nucleation site at the enhancer (2). Phosphorylation of RNAPII-CTD on serine 2 (3) leads to disassembly of the NANOG-RNAPII complex, resulting in eRNA synthesis at the enhancer and release of locally high concentrations of RNAPII in the vicinity of nearby promoters.

### Phosphorylation of RNAPII CTD serine-2 breaks the NANOG complex

RNAPII is phosphorylated on serine-2 by CDK9 (*25*). To determine whether RNAPII within the RNAPII:NANOG complex is a substrate for CDK9, GST-CTD:NANOG complex was first purified on GSH-resin. Subsequent treatment with CDK9 showed that RNAPII within the RNAPII:NANOG complex could be phosphorylated by CDK9 (Fig. 4b). More importantly, NANOG was dissociated from the RNAPII:NANOG complex by CDK9 in an enzyme- and ATP-dependent manner (Fig. 4b). Together, the preceding results show that NANOG can bind to unmodified RNAPII and to phospho-serine 5 modified RNAPII both of which are present before and during transcriptional pausing (*26*), but that the enzyme driving pause release causes NANOG-RNAPII complex dissociation.

To determine whether CDK9 also affects the RNAPII-NANOG complex in cells, ESCs were cultured in the presence of the CDK9 inhibitor, flavopiridol. CUT&RUN showed that flavopiridol treatment resulted in accumulation of NANOG on chromatin (Fig. 4c, Supplementary data Fig. 7). This accumulation was specific for NANOG binding regions that were co-bound by RNAPII as no increase was seen at NANOG binding sites not associated with RNAPII. Moreover, this accumulation was more pronounced at sites associated with NANOG DEGs (Fig. 4c). Together these data suggest a mechanism by which NANOG-RNAPII binding on chromatin can be dissociated by a known regulator of transcription.

## Discussion

Models of mammalian transcription classically propose that TFs bind DNA and that intermediary proteins connect TFs to the transcriptional machinery (*27*). While recent studies have introduced the potential of molecular condensates to regulate enhancer function and control transcription, current thinking retains a focus on the proposed indirect nature of connections between TFs and RNAPII (*28, 29*). In contrast, our study provides the first example in mammalian cells of direct contact between a cell-type specific DNA binding TF, NANOG and RNAPII. How generalisable might this finding be? Possibly, the connection between NANOG and RNAPII is unique. Alternatively, the high density of tryptophan residues (10 within 46 residues) in the WR may have sensitized detection of a more widespread phenomenon. As we propose that NANOG-RNAPII binding is mediated by aromatic stacking, it is noteworthy that the extended aromaticity of the indole ring of tryptophan strengthens the π-π interactions responsible for aromatic stacking compared to the benzyl rings of phenylalanine or tyrosine (*30*). This may have sufficed to bring RNAP2 above the detection threshold in the interactome of this TF (*11*). A prominent role for tryptophan residues in transcriptional activation has also been highlighted in a functional screen (*31*).

Our demonstration that a sequence specific DNA binding TF connects DNA directly to RNAPII has implications for how such TFs can operate. Interestingly, p53, a ubiquitously expressed sequence specific DNA binding TF, has also been shown to bind directly to RNAPII (*32, 33*). However, there are differences between binding of p53 and NANOG to RNAPII. p53 binds RNAPII through interactions of the core DNA binding domain with the RNAPII DNA binding cleft where the Rbp1 clamp meets upstream DNA, and through interactions of the p53 transactivation domains with the RNAPII jaw, where Rbp1 meets downstream DNA (*33*). In contrast, NANOG binds to the CTD of RNAPII subunit Rbp1, which is distal to the DNA binding site (*34*). While Nanog and p53 bind RNAPII in disparate ways, they both bind predominantly to enhancers (*35, 36* and data shown here). How these different mechanisms of RNAPII interaction influence enhancer function is currently unclear. However, in the context of NANOG function, it is interesting that the RNAPII CTD has been reported to be required for enhancer activity (*37*). In addition, shortening the CTD to 25 repeats in human cells reduces enhancer transcription and transcriptional activation in response to exogenous signals but has a limited effect on steady state transcription (*38*). Inhibition of CDK9 by flavopiridol is known to increase the level of RNAPII bound to chromatin and has been reported to inhibit pause-release of RNAPII at both promoters and enhancers (*39–42*). Flavopiridol also stimulated an increase in chromatin binding by NANOG, suggesting that the NANOG:RNAPII complex interacts with chromatin. This idea is supported by both NANOG SICAP (*43*), and directly by our demonstration that NANOG recruits RNAPII to DNA and chromatin in a WR-dependent manner. Therefore, RNAPII may associate with ESC enhancers more frequently when NANOG is bound to them. Furthermore, the affinity with which NANOG binds DNA is enhanced by RNAPII CTD, suggesting that NANOG may bind more strongly to chromatin microenvironments where the RNAPII concentration is increased. In this regard it is notable that the yeast RNAPII CTD has recently been shown to constrain RNAPII in space, effectively increasing the local RNAPII concentration (*44*). In addition, the ability of CDK9 to disrupt the RNAPII-NANOG complex may suggest that localized CDK9 activity could result in a localized reduction in the avidity of NANOG for chromatin.

Our findings suggest that the interaction with NANOG can concentrate RNAPII at enhancers. This may facilitate transfer of RNAPII to the relevant NANOG-responsive promoter. While this could occur through one of several classic models (*27*), the fact that both RNAP2 and NANOG have been reported to undergo phase separation events (*45–49*), indicates the possibility that indirect connections mediated by phase separated condensates could be involved (*28, 29*). Alternatively, transfer may occur independent of condensate formation by controlled aggregation from a nucleation site at an enhancer defined by binding of NANOG (Fig. 4d). As NANOG and RNAPII interact using tryptophans and tyrosines present multiple times within their low complexity domains, binding of RNAPII to NANOG could then initiate a series of NANOG-RNAPII interactions by using more than one site on each multivalent monomer (*50*). Reiterative recruitment could then synthesise a domain of increasing size extending in space from the enhancer. Synthesis of a domain of increasing size from a nucleation site at an enhancer could also explain how the *Snail* shadow enhancer can drive simultaneous transcriptional bursts from two separate, but equidistant promoters (*51*). A further intriguing possibility suggested by the observation that CDK9 dissociates the NANOG:RNAPII complex is that CDK9 may act on a domain extending in space from an enhancer, potentially resulting in eRNA transcription and release of RNAPII in the vicinity of a responsive promoter.

## Materials and Methods

### Co-immunoprecipitation

Co-immunoprecipitations were performed as described (*13*) (*13*). E14/T cells were transfected with pPyCAG(Flag)_3_-Nanog IP and nuclear lysates prepared 24 hours post-transfection. Immunoprecipitations were performed with antibodies raised to RNAPII and co-precipitating NANOG was detected by immunoblotting. Antibodies used were 3D12 (phospho-Y1), H5 (phospho-S2) and CTD4H8 (phospho-S5).

### Recombinant protein

Recombinant full length NANOG was expressed in *E. coli* from a pET15-Nanog construct. Cells were induced with 1mM IPTG once cell density reached an O.D. of ~0.6 and grown at 37°C for a further 3 hours. Inclusion bodies were prepared by sonication and centrifugation and then solubilized in 25mM Hepes pH7.6, 7M urea, 200mM NaCl, 10mM β-mercaptoethanol, 10mM imidazole. Protein was purified on nickel resin (Cat. No. P6611, Sigma) and NANOG containing fractions pooled. Protein was concentrated to 10mg/ml in a Vivaspin device with a molecular weight cut off of 10kDa (Sartorious, Cat. No. VS2002). Soluble, refolded material was obtained by rapid 10-fold dilution into 10mM Hepes pH7.9, 50mM KCl, 10mM NaCl, 10% glycerol, 2.5mM DTT, 0.5mM EDTA.

Recombinant CTD was expressed as a GST-fusion in pGEX-2T. Cells were induced with 1mM IPTG once cell density reached an O.D. of ~0.6 and grown at 18°C for a further 5 hours. Cells were sonicated in PBS and protein purified from the soluble lysate on GSH-sepharose resin (Cytiva, Cat. No 17075601).

### Recombinant protein co-expression

GST-CTD or MBP-WR were co-expressed with NANOG in *E. coli* BL21(DE3) from pET21, pMAL and pRSF plasmids respectively. Expression was induced by 1mM IPTG at 18°C for 5 hours. Cell lysates were prepared by sonication. Purification of NANOG was performed on nickel resin (cat. no. P6611, Sigma) and MBP tagged constructs were purified on amylose resin (cat. no. E8021, NEB). For CDK9 treatments, protein bound beads were washed in 25mM Tris pH7.6, 10mM MgCl_2_, 5mM β-glycerophosphate, 2.5mM DTT and 0.2μg CDK9 (cat. no PV4131, ThermoFisher) added in a final volume of 50μl. Reactions were incubated (30°C, 1 hour) before further washing and elution of bound material in Laemmli buffer.

### Electrophoretic shift mobility assays

EMSAs were performed in 10mM Hepes pH7.9, 50mM KCl, 10mM NaCl, 10% glycerol, 2.5mM DTT, 0.5mM EDTA and 0.1mg/ml BSA. The probe used is a 26mer oligonucleotide probe derived from the *Tcf3* promoter (*17*) labelled with IRDye 680. Mixes containing oligonucleotide (5nM) and protein (30nM) were incubated (30mins, room temperature) prior to loading onto gels. For K_D_ measurements the oligonucleotide concentration was 5nM with NANOG titrated down from 500nM (0.66 fold steps) in the presence or absence of 1.5μM GST-CTD.

### BASIL assay

Streptavidin beads (Thermo, Cat. No. 20347) were pre-blocked with chicken egg albumin at 200ng/μl for 1 hour. Beads were then loaded with biotinylated double stranded oligonucleotide at 1.5μM for 1 hr. The oligonucleotide sequence is the same as used for EMSA. After washing 3x with PBS, beads were incubated with rNANOG at 1mg/ml or with for 1 hour. After washing 3x with PBS, beads were incubated with fluorescein amidite labelled oligonucleotide (unbiotinylated) of the same sequence described above, for 1 hour. After washing 3x with PBS beads were visualized.

### RNA analysis

RNA analysis was performed as described in Festuccia et al(*10*)

### Chromatin immunoprecipitation

10 million ESCs were plated and cultured for 16 hours before crosslinking on plate with 1% formaldehyde (Sigma) (10 min, 37°C). The reaction was quenched with 0.125 mM glycine (5 min, room temperature). Cells were washed three times with PBS (4°C). Cells were covered with swelling buffer (25mM Hepes pH7.9, 1.5mM MgCl2, 10mM KCl, 0.1% NP-40) and incubated (20 mins, 4°C). Nuclei were harvested by scraping and resuspended in sonication buffer (50mM Hepes pH7.9, 140mM NaCl, 1mM EDTA, 1% Triton X100, 0.1% deoxycholate, 0.1% SDS). Samples were sonicated on ice in a Bioruptor Pico (Diagenode). Immunoprecipitations were performed using 5μg of antibody (RNAPII (SC-899), NANOG(*52*), by rotation overnight at 4°C, in a final volume of 1ml. Immunocomplexes were purified using Protein-G Dynabeads (Lifetech). Libraries were prepared using the NEBNext Ultra II kit (E7645). For flavopiridol experiments, inhibitor was added to a final concentration of 10μM for the times indicated.

### CUT&RUN

Cleavage Under Targets & Release Using Nuclease (CUT&RUN) was performed as previously reported with no modifications using 5×10^5^ cells per replicate, 2 biological replicates per sample, 1:100 dilution of anti-NANOG REF and homemade pAG-MNase. Barcoded libraries were generated with the NEBNext Ultra II DNA Library prep kit (E7645S) and NEBNext Multiplex Oligos for Illumina (E7335S, E7500S, E7710S, E7730S) with modifications. 5 ng of DNA was used per library. End-prep and ligation were performed according to the manufacturer’s instructions. 1 µl of USER enzyme was used per sample. For the final PCR step, 1 µl of both universal and indexed primers were used, and the annealing/extension step was decreased to 30 seconds to reduced amplification of large fragments. Clean up steps were performed with home-made PCR purification beads. Library quality was assessed with HSD1000 ScreenTapes (Agilent, cat. 5067-5584 and 5067-5585) in a Tapestation 2200 (Agilent). Libraries were sequenced paired-end on Illumina NextSeq 500 and NextSeq 2000.

### Bioinformatic analysis

For ChIP-seq and CUT&RUN the quality of reads was assessed by FastQC (v. 0.11.9), index and adaptor sequences were trimmed using TrimGalore (v. 0.6.6) and Cutadapt (v. 1.9.1) followed by a second quality check with FastQC. Sequences were aligned to the mm10 mouse reference genome using Burrows-Wheeler Alignment (BWA, v. 0.7.16) tool with the maximal exact matches (MEM) option. Fragments were quality filtered (>10) with Samtools (v. 1.6) and PCR duplicates were marked with Picard (v. 2.23.3) and removed with Samtools. CUT&RUN reads in ESCs were normalized to the number of *E. coli* reads before further analysis.

Peaks were called with Model-Based Analysis of ChIP-Seq (MACS2, v. 2.1.1) at 5% FDR using parameter -f BAMPE for paired-end input. Blacklisted regions were subtracted from the peak list using BEDTools (v. 2.27.1). Bigwig tracks were generated with the bamCoverage option in the python-based DeepTools (v. 3.5.1).

## ACKNOWLEDGEMENTS

We thank G. Hateminejad for technical assistance and A. Soufi, S. Lowell, S. Pollard and D. O’Carroll for comments on the manuscript. This work was supported by grants to IC from the Medical Research Council (MR/T003162/1) and the Biotechnology and Biological Sciences Research Council grant (BB/T008644/1).

## AUTHOR CONTRIBUTIONS

NPM and IPC conceived the project and designed experiments. NPM performed the majority of the experiments and with IC interpreted the data. EB performed CUT&RUN analyses and performed bioinformatics analyses, JZ performed RNA expression analysis and with MV performed ChIP experiments. AMcG and ST performed bioinformatics analyses. DC performed cell culture and molecular biology experiments. IC and NM wrote the manuscript.

---

**Supplementary data figure S1.**
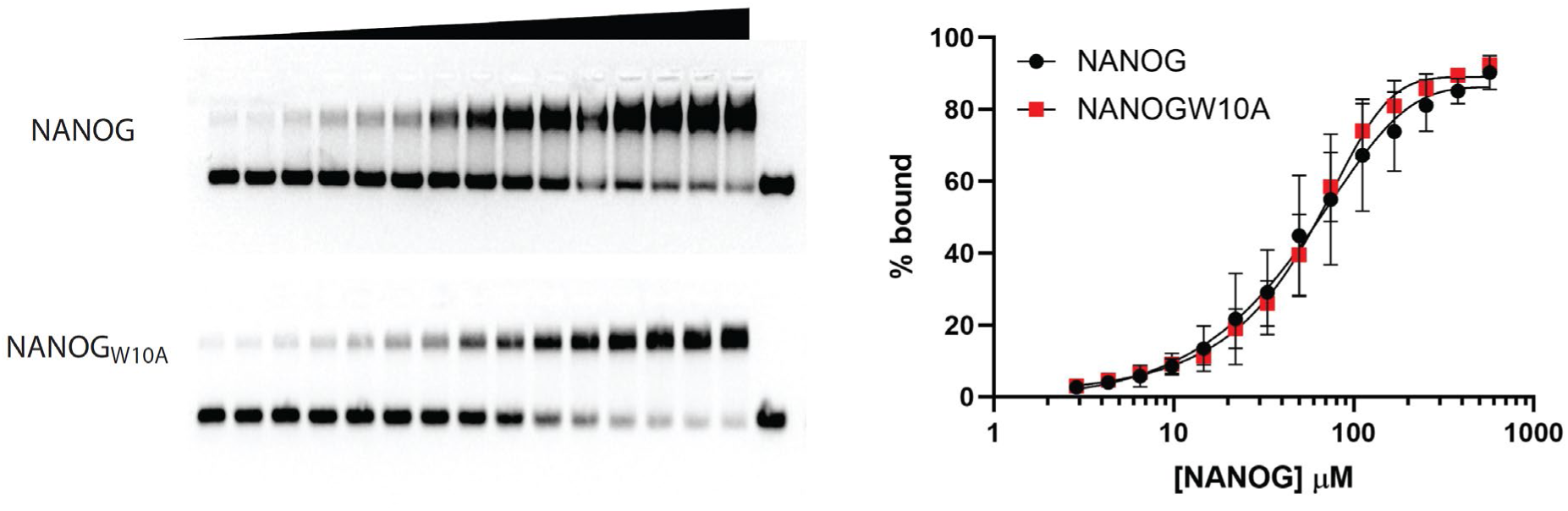
Electrophoretic mobility assays with NANOG and NANOG_W10A_. EMSAs were performed with titrations of NANOG and NANOG_W10A_. Proteins were titrated from 500nM in a 1/3 dilution series (n=3, error bars represent standard deviation).

**Supplementary data figure S2.**
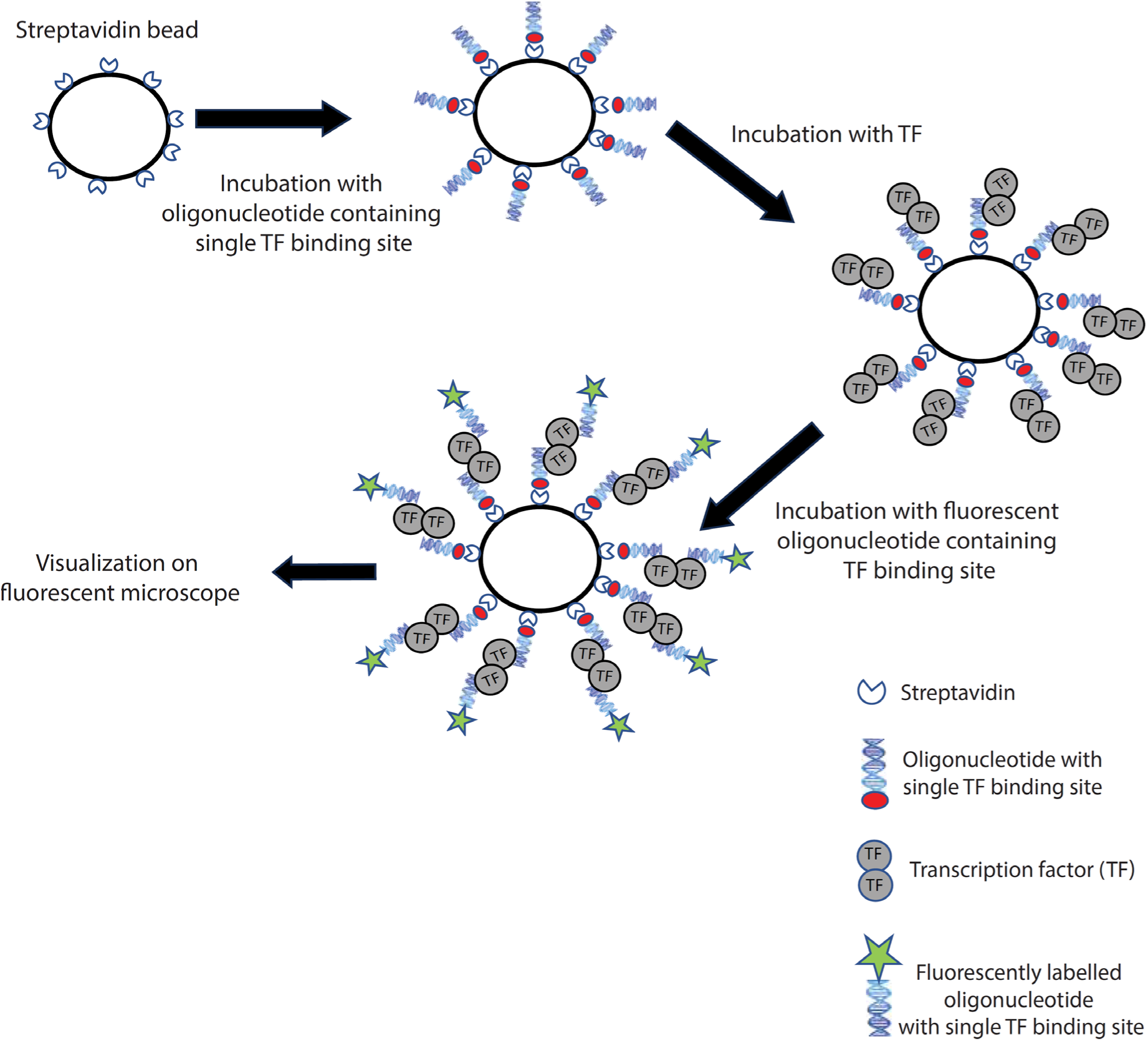
Illustration of BASIL bead binding assay. If a DNA bound TF can form homo-multimers, this presents a second DNA binding domain that is accessible for binding a fluorescently labelled oligonucleotide.

**Supplementary data figure S3.**
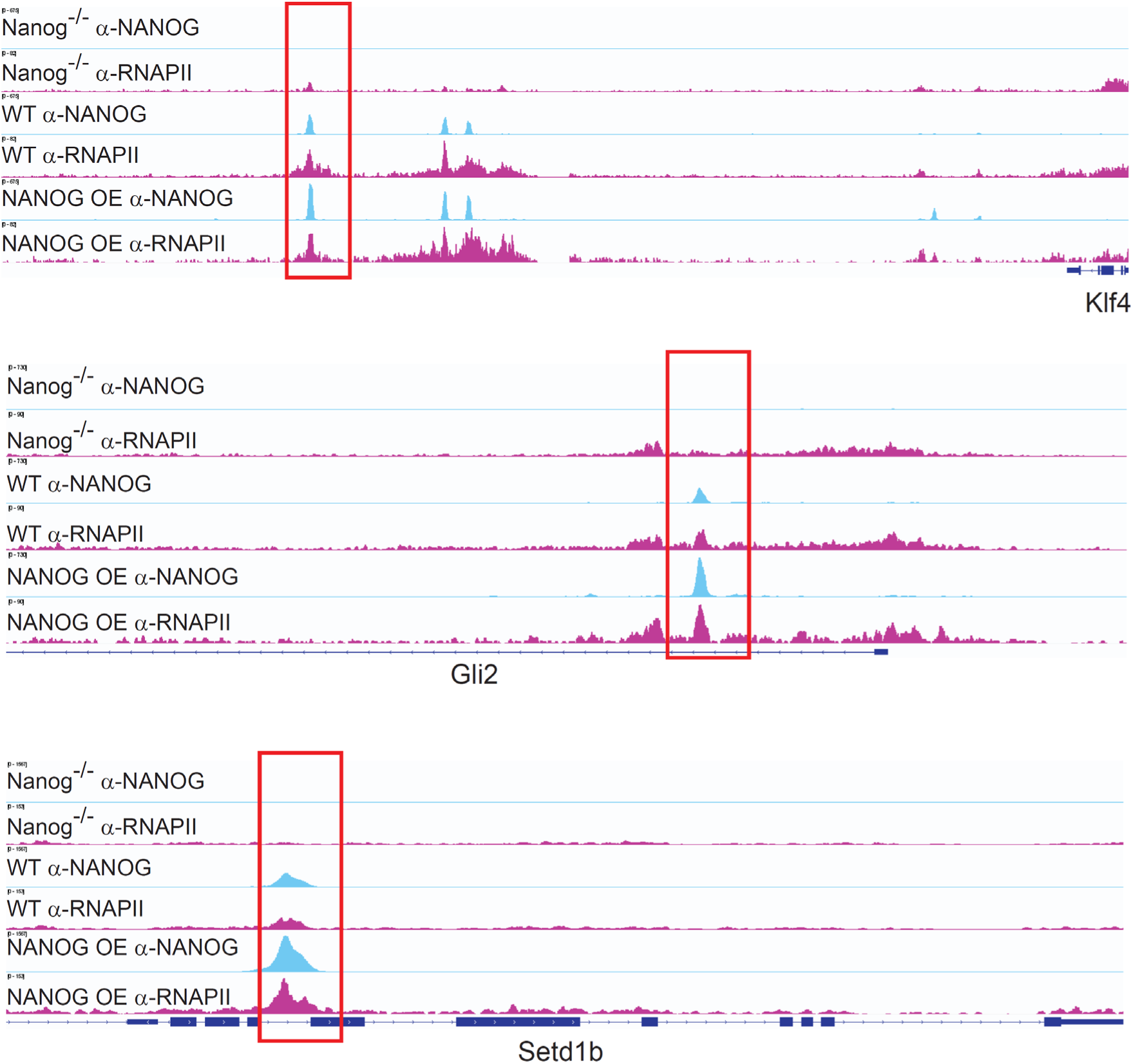
ChIP-seq in *Nanog*^−/−^, WT and NANOG over-expressing ESCs. ChIP-seq tracks for NANOG and RNAPII around *Klf4* (top), *Gli2* (middle) and *Setd1b* (bottom). The red boxes highlight where increased NANOG signal results in increased RNAPII binding.

**Supplementary data figure S4.**
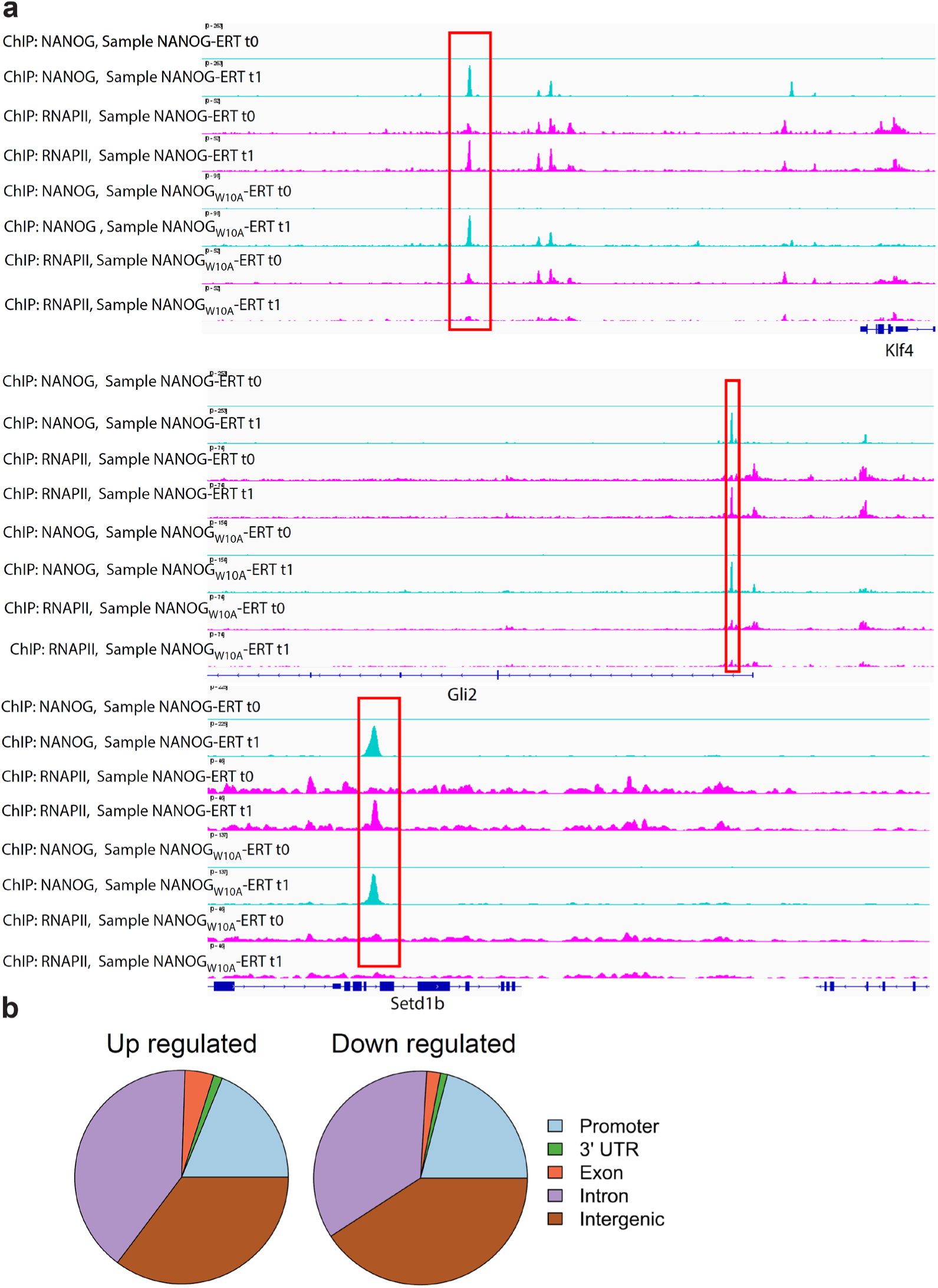
ChIP-seq analysis of NANOG-ERT^2^ and NANOG_W10A_–ERT^2^. **a.** Examples of ChIP-seq tracks for NANOG-ERT^2^, NANOG_W10A_–ERT^2^ and RNAPII around genes upregulated by NANOG 0hr (t0) and 1hr (t1) after addition of tamoxifen. Tracks are *Klf4* (top), *Gli2* (middle) and *Setd1b* (bottom). The red boxes highlight the RNAPII peaks increased by the presence of NANOG. b. Venn diagrams of NANOG occupancy at NANOG responsive genes.

**Supplementary data figure S5.**
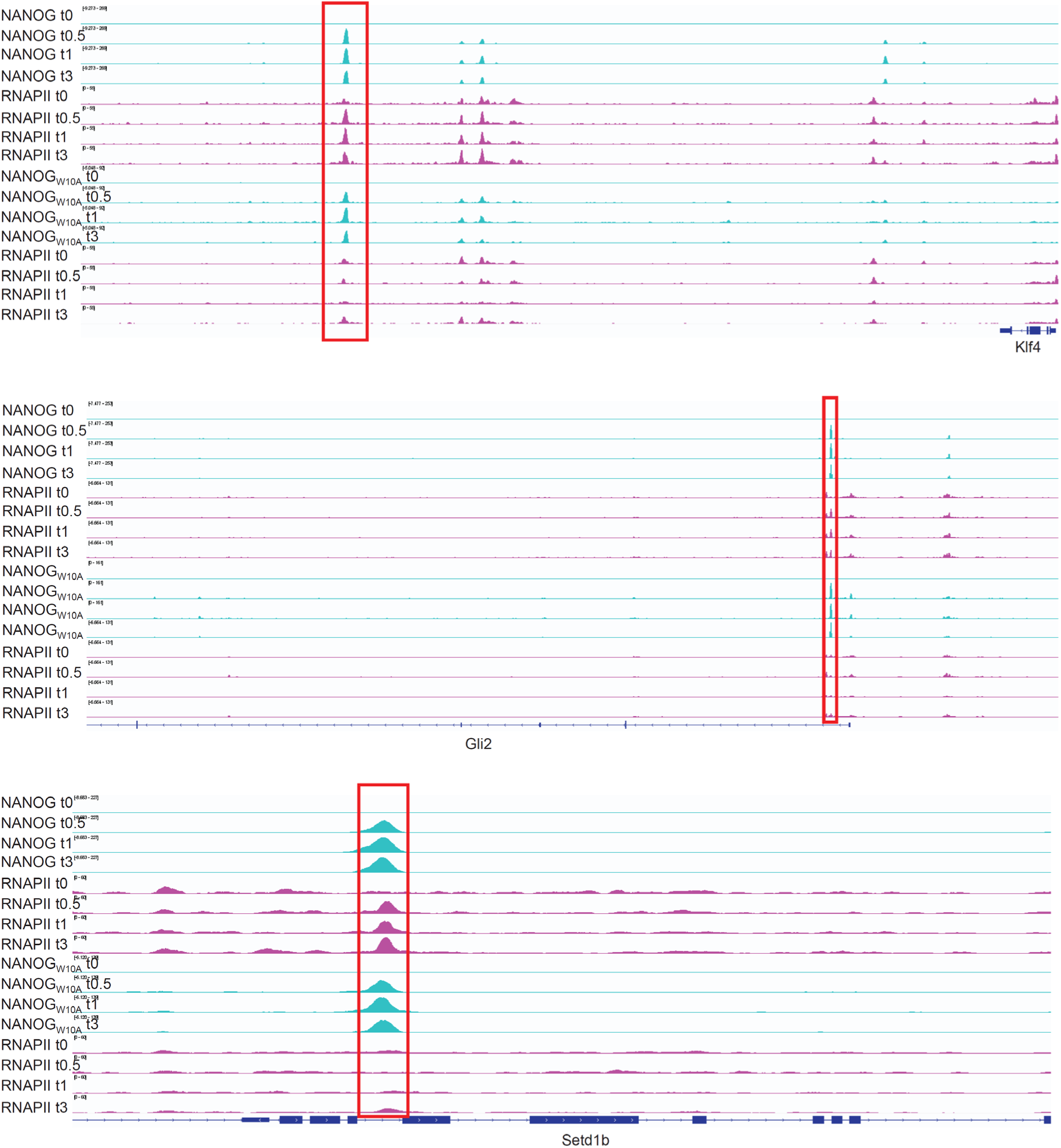
ChIP-seq analysis of time course of induction of NANOG-ERT^2^ and NANOG_W10A_–ERT^2^. **a**. Examples of ChIP-seq tracks for NANOG-ERT^2^, NANOG_W10A_–ERT^2^ and RNAPII around genes upregulated by NANOG 0hr (t0), 0.5hr (t0.5), 1hr (t1) and 3hr (t3) after addition of tamoxifen. Tracks are *Klf4* (top), *Gli2* (middle) and *Setd1b* (bottom). The red boxes highlight the RNAPII peaks increased by the presence of NANOG.

**Supplementary data figure S6.**
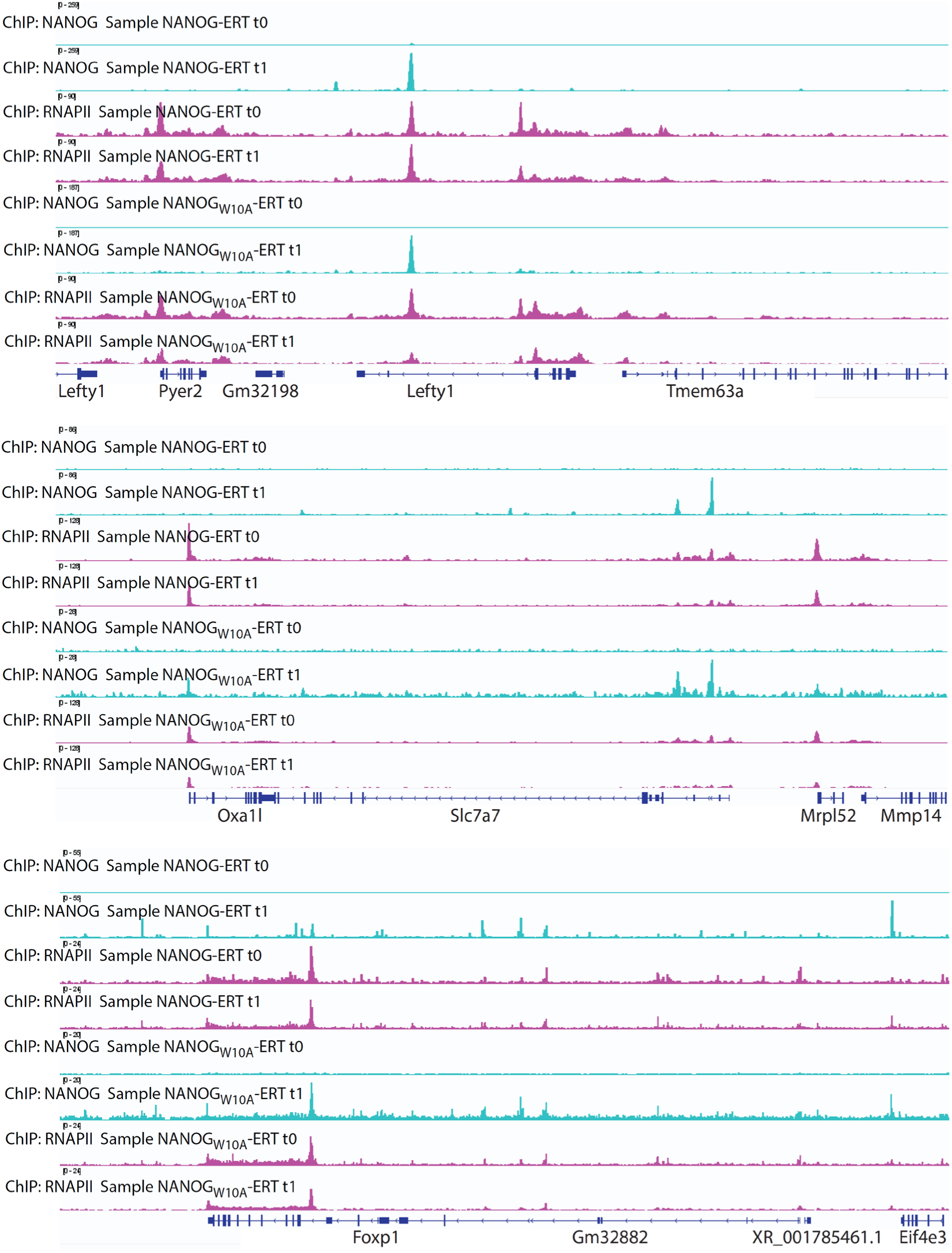
ChIP-seq analysis of NANOG-ERT^2^ and NANOG_W10A_–ERT^2^. Examples of ChIP-seq tracks for Nanog, W10A and RNAPII around genes downregulated by NANOG 0hr (t0) and 1hr (t1) after addition of tamoxifen. Tracks are *Lefty1* (top), *Slc7a7* (middle) and *Foxp1* (bottom). The red boxes highlight the regions where NANOG and RNAPII peaks overlap.

**Supplementary data figure S7.**
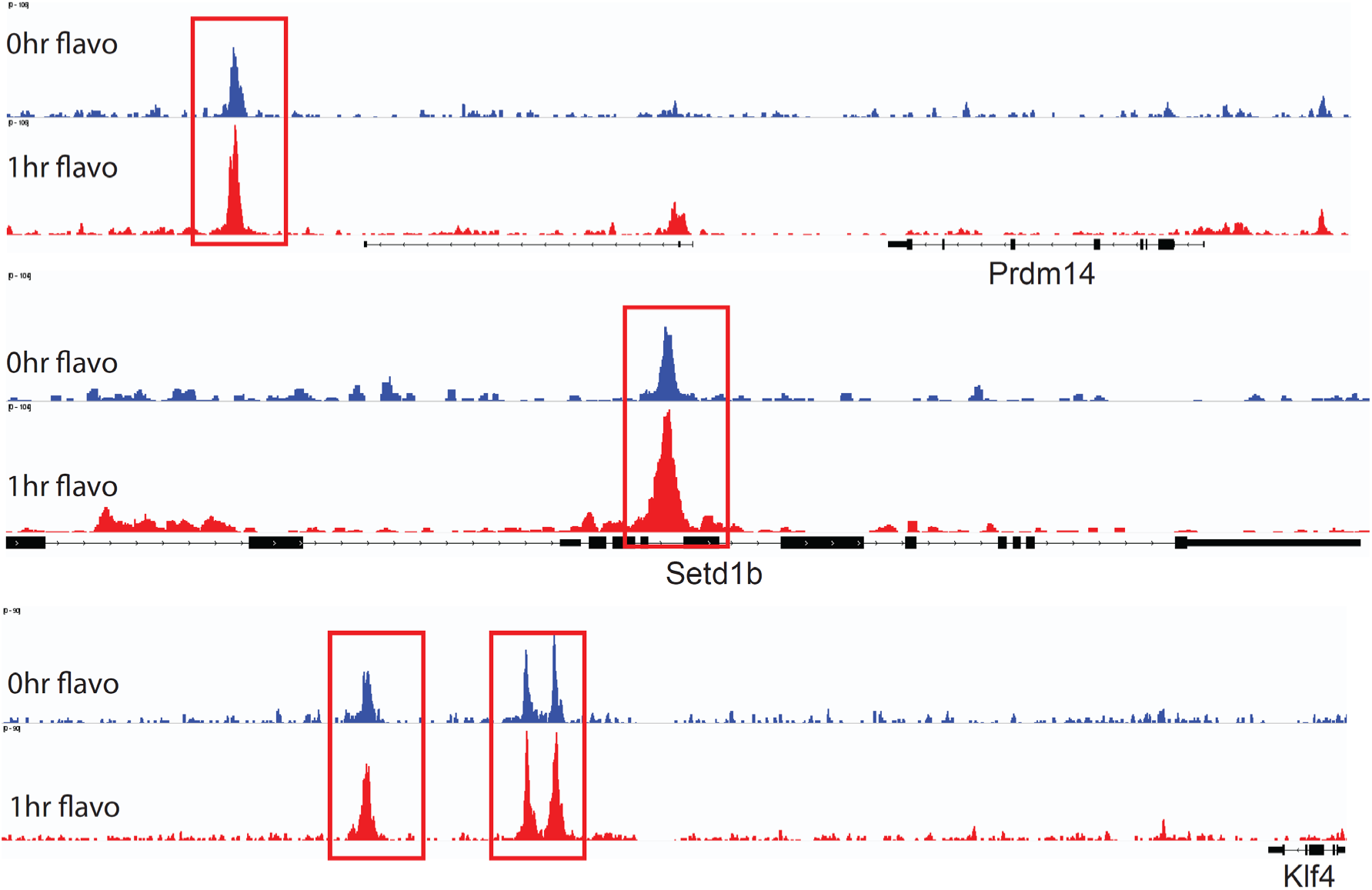
ChIP-seq tracks of NANOG binding in WT ESCs treated with flavopiridol. Tracks from untreated cells (blue) or after 1 hour of flavopiridol treatment (red). Tracks are *Prdm14*, *Setd1b*, *Klf4*.

**Supplementary table S1.**
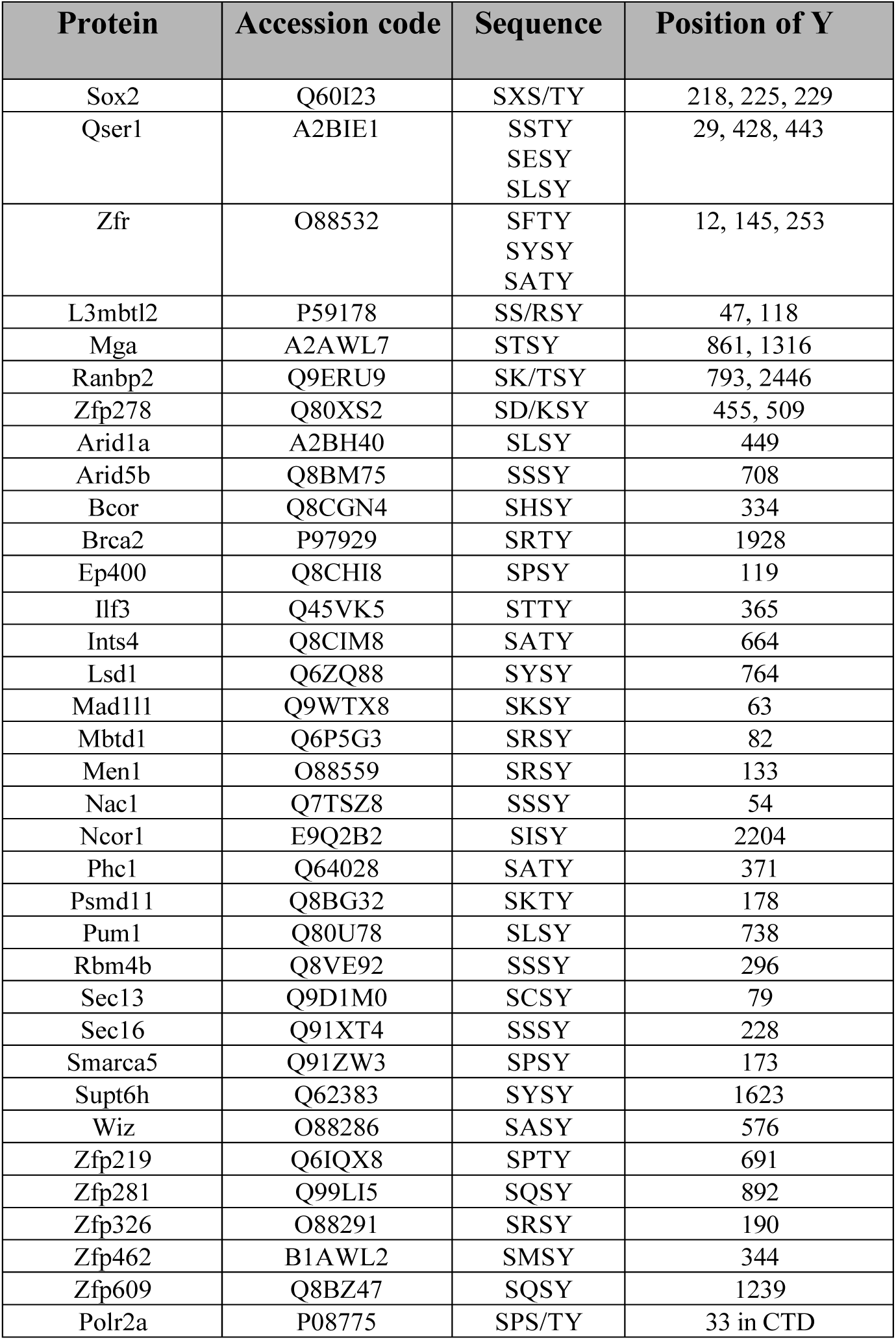
Proteins containing the SXS/TY motif identified as the NANOG binding sequence in Sox2 (*11*). The sequence and the position of the motif(s) within the protein are shown.

